# Dual RNA sequencing (dRNA-Seq) of bacteria and their host cells

**DOI:** 10.1101/098715

**Authors:** James W. Marsh, Regan Hayward, Amol Shetty, Anup Mahurkar, Michael S. Humphrys, Garry S. A. Myers

**Affiliations:** iThree Institute, University of Technology, Sydney, New South Wales, Australia JM: RH: GM; Institute for Genome Sciences, University of Maryland School of Medicine, Baltimore, Maryland, United States of America AS: AM: MH

## Abstract

Bacterial pathogens subvert host cells by manipulating cellular pathways for survival and replication; in turn, host cells respond to the invading pathogen through cascading changes in gene expression. Deciphering these complex temporal and spatial dynamics to identify novel bacterial virulence factors or host response pathways is crucial for improved diagnostics and therapeutics. Dual RNA sequencing (dRNA-Seq) has recently been developed to simultaneously capture host and bacterial transcriptomes from an infected cell. This approach builds on the high sensitivity and resolution of RNA-Seq technology and is applicable to any bacteria that interact with eukaryotic cells, encompassing parasitic, commensal or mutualistic lifestyles. We pioneered dRNA-Seq to simultaneously capture prokaryotic and eukaryotic expression profiles of cells infected with bacteria, using *in vitro Chlamydia*-infected epithelial cells as proof of principle. Here we provide a detailed laboratory and bioinformatics protocol for dRNA-seq that is readily adaptable to any host-bacteria system of interest.

## Introduction

### Background

Upon infection or other interactions, bacteria and their host eukaryotic cells engage in a complex interplay as they negotiate their respective survival and defense strategies. Unraveling these coordinated regulatory interactions, virulence mechanisms, and innate responses is key for our understanding of pathogenesis, disease and the development of therapeutics [1]. Traditional transcriptomic approaches such as microarrays have typically focused on *either* the prokaryotic or eukaryotic organism to investigate the host-bacteria interaction network [2]. However, this approach cannot decipher reciprocal changes in gene expression that contribute to the global infection system. Instead, an integrated approach is required that acknowledges both interaction partners, i.e. both bacteria and host, from the same biological sample. Due to the increasing affordability and resolution of next-generation sequencing, this is now achievable via dual RNA sequencing (dRNA-Seq) [1].

RNA-Seq was developed for the study of transcriptomes based on the massively parallel sequencing of RNA [3]. In a typical experiment, total mRNA from a sample is subjected to high-throughput next-generation sequencing and mapped to a reference genome to deduce the structure and/or expression state of each transcript [4]. Gene expression changes can be accurately measured between samples with high coverage and sensitivity, while alternative splicing analyses can be applied to identify novel isoforms and transcripts, RNA editing, and allele-specific expression [5]. The high sensitivity and dynamic range of RNA-Seq has expanded our capability for whole transcriptome analysis and enabled new insight into the functional elements of the genome [6].

dRNA-Seq extends these capabilities to two (or potentially more) interacting organisms, allowing the simultaneous monitoring of gene expression changes without disturbing the complex interactions that define host-bacteria infection dynamics. We applied dRNA-Seq to map host and bacteria transcriptomes from *Chlamydia-infected* host epithelial cells, which highlighted a dramatic early response to infection and numerous altered pathways within the host cell [1]. dRNA-Seq has since been successfully used to study host-bacteria interactions for *Salmonella enterica* [7], *Azospirillum brasilense* [8], *Mycobacterium tuberculosis* [9], *Haemophilus influenzae* [10], *Yersinia pseudotuberculosis* [11] and *Actinobacillus pleuropnemoniae* [12].

### Advantages and limitations

cDNA microarrays first enabled large-scale transcriptome analyses, allowing the expression pattern of tens of thousands of known genes to be measured. Drawbacks include (1) a high background signal [13]; (2) cross-hybridization between genes of similar sequence; (3) the limit of expression level detection to the 1,000-fold range, compared to the actual cellular 1,000,000-fold range [14] (4) restriction of analysis to known or predicted mRNAs [15]; and (5) the inability to detect novel transcripts [14]. Some of these were overcome with tiling arrays to measure antisense RNA expression and other non-coding RNA (ncRNA) transcripts, but the large size of eukaryotic genomes make this inordinately costly [16]. Tag-based sequencing does enable the enumeration of individual transcripts, but this method requires existing gene structure information, can only sample a small region of a transcript, and is incapable of capturing diverse classes of RNA and its isoforms.

RNA-Seq provides a wider dynamic range, higher technical reproducibility, and a better estimate of absolute expression levels with lower background noise [17-19], and has become the primary method to examine transcriptomes. By allowing an unbiased determination of gene expression, high resolution data on potentially transcribed regions upstream and downstream of the annotated coding region, and post-translational rearrangements such as splicing and different RNA isoforms can be reported [20]. As a result, RNA-Seq improves genome annotation, and identifies new ORFs, transcription start sites (TSSs), the 5’ and 3’ UTRs of known genes, non-coding RNAs such as microRNA, promoter-associated RNA, and antisense 3’ termini-associated RNA [21]. dRNA-Seq can report these data for two (or potentially more) organisms from the same sample, while providing powerful insight into novel interaction dynamics. For example, gene expression changes in one organism can be correlated with the responses of the other to capture crucial events that signify the dynamic mechanisms of host adaption and the progression of infection [1,4,7,10,22].

Despite these advantages, dRNA-Seq remains technically challenging. Up to 98% of total RNA is ribosomal (rRNA) [23], while bacterial mRNA levels are typically low compared to the host, especially during early infection periods, requiring mRNA depletion and/or enrichment approaches for cost-effective sequencing. Additionally, the quantity of mRNA detected by RNA-Seq is often a poor indicator for protein abundance due to mRNA instability and turnover [24,25]. The wide range of expression levels can result in non-uniform coverage where only a few reads can be captured for genes subject to lower expression levels, while short isoforms and repeat sequences derived from the same gene may result in assembly ambiguities. These ambiguities are compounded when using *de novo* methods for genomes that are partially or fully un-sequenced [17], but can be avoided when assembling reads to a reference genome. Transcript length bias can distort the identification of differentially expressed genes in favor of longer transcripts [26], but can be standardized with appropriate normalization techniques. Despite these challenges, dRNA-Seq is a powerful, economical, sensitive, and species-independent platform for investigating the gene expression dynamics of host-bacteria interactions [4].

### Overview of the technique

This protocol provides a clear and practical laboratory and a detailed bioinformatics analysis pipeline for a typical dRNA-Seq host-bacteria analysis. We describe an experiment based on human epithelial carcinoma (HeLa) cells (host) infected with *Chlamydia trachomatis* serovar E (bacteria), which is readily adaptable to any host-bacteria system of interest.

*C. trachomatis* is an obligate intracellular bacterial bacteria that is reliant on its host cell for survival. It has been otherwise recalcitrant to standard genetic analyses, rendering it ideal for the development and application of dRNA-Seq. The protocol includes all steps for the bacterial infection of host cells, total RNA extraction, and rRNA depletion in preparation for sequencing (Figure 1). Bioinformatic steps are then included for total RNA sequence quality control and trimming, the *in silico* segregation of host and bacteria reads, distinct sequence alignment and sorting techniques for host and bacteria data, alignment visualization, read quantification and normalization, the separate statistical analysis of host and bacteria data, and the final integration of host and bacteria transcriptomic responses (Figure 2).

**Figure 1.**
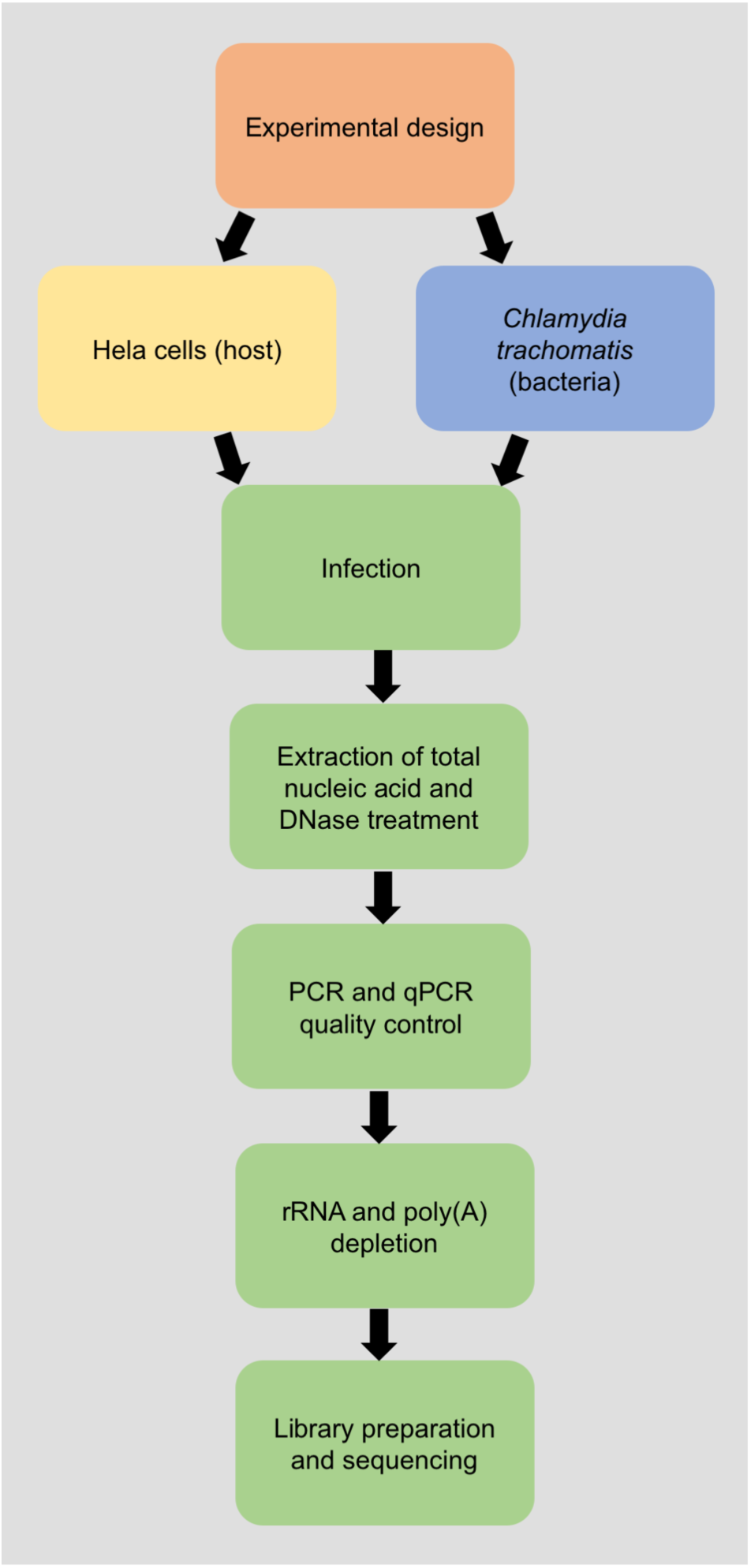
Flow-chart for the wet-lab protocol for dRNA-seq of bacteria and host.

**Figure 2.**
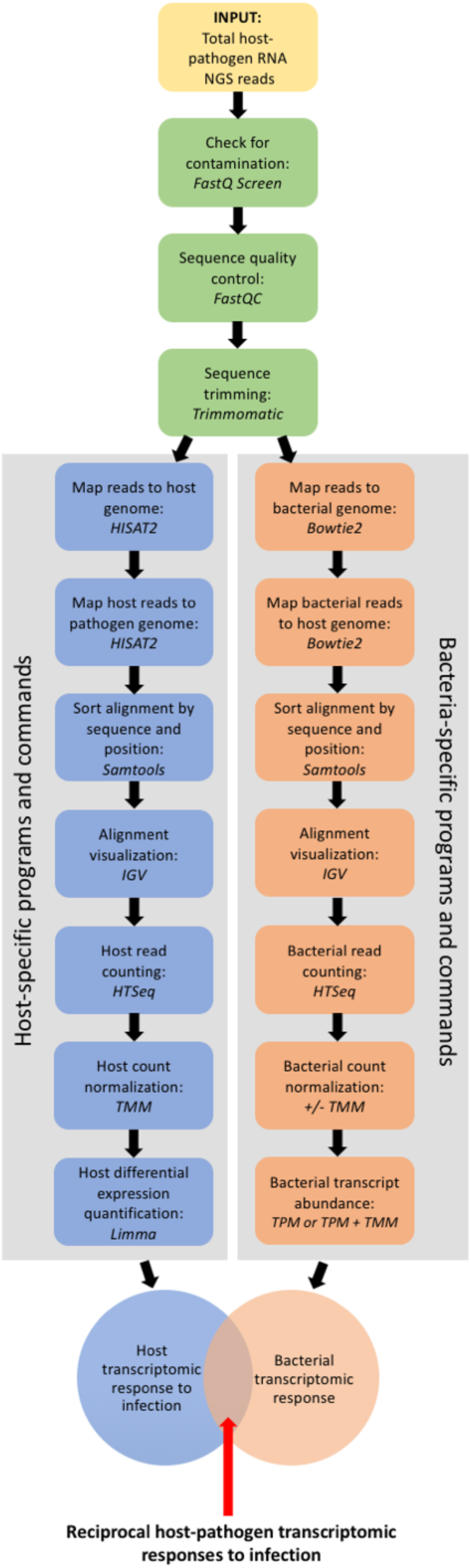
Flow-chart for the bioinformatic data analysis of dRNA-Seq of host and bacteria.

From here, the user may diverge toward a number of possible downstream applications, including time-series analysis, host alternative splicing in response to the bacteria, gene set enrichment and ontology analysis, and elucidation of interaction and regulatory networks [27]. dRNA-Seq data may be integrated with other data sources to establish a more complete picture of gene regulation, including genotyping data to identify genetic loci responsible for gene expression variation, epigenetic information to highlight the influence of transcription factor binding, histone modification, and methylation, and miRNA-Seq data to identify the regulatory mechanisms of gene expression changes via non-coding RNA [28,29].

### Experimental design

For RNA-Seq, a typical workflow includes experimental design, RNA extraction, library preparation, sequencing, and data analysis. The experiment should be designed to address the biological question(s) of interest and a key starting point is to determine the RNA species to be investigated (i.e. mRNA, miRNA, snRNA etc). This will influence the quantity of input RNA and sequencing depth required, which are both crucial factors for successful dRNA-Seq experiments. To capture sufficient RNA from both organisms, the ratio of bacteria to host genome size is a useful starting point followed by an estimation of the desired fold coverage. This can be determined by considering the number of replicates, the expected influence of housekeeping and structural RNA (rRNA and tRNA), the possibility of host:bacteria sequence overlap, the number of time-points, and the multiplicity of infection (MOI). We suggest at least three biological replicates for each sample rather than technical replicates taken from the same sample to minimize Type I and II errors and ensure an adequate estimation of within and between sample variation. As ∼95% of total RNA will be ribosomal, a method of rRNA depletion and/or mRNA enrichment is recommended and several options are discussed below. Sequence overlap between host and bacteria can be predicted by mapping the bacterial sequence reads to the host genome and vice versa, which is also discussed below. The time-points of interest should be carefully considered as the initiation and period of transcriptional response can differ between host and bacteria [30]. Ideally, multiple time-points should be collected to suitably capture the dynamic host-bacteria transcriptional landscape. Finally, a suitable bacteria MOI should be selected to maximize the transcriptional signal from both host and bacteria, while reducing bias towards the uninfected cells that will flourish at the later time-points. Importantly, a high(er) MOI may also lead to a heightened and/or distorted host response with decreased biological relevance, depending on the system under investigation. Deeper sequencing may be necessary for the detection of low copy number transcripts or alternate isoforms, however increased sequencing depth can also increase the detection of transcriptional noise, spurious cDNA transcripts, or genomic DNA contamination so careful consideration is required [31,32]. Optionally, the addition of RNA spike-ins and unique molecular identifiers (UMI) can be useful for the quantitative calibration of RNA levels [33,34].

This protocol describes the collection and analysis of protein-coding mRNA from *C. trachomatis* and Hela cells at 1 and 24 hours post-infection (hpi) time-points as they are physiologically relevant to the early and mid-stages of the chlamydial development cycle. An MOI of 1.0 is used to maximize chlamydial RNA recovery, however a limited quantity of bacterial RNA will be present at 1 hpi as replication would not yet have commenced. This caveat should be considered when estimating RNA quantities required. It is recommended that ∼ 5 × 10^8^ host reads and > 1 × 10^6^ bacterial reads are required for adequate coverage [4,35]. The *Chlamydia* to Hela genome size is ∼1:3200 MB, indicating that *Chlamydia's* RNA accounts for ∼0.03% of total host-bacteria RNA. As ∼95% of this will be uninformative rRNA and tRNA [4], 0.0015% and 4.9985% of total RNA will represent informative bacteria and host mRNA, respectively. Given this ratio, 1 × 10^10^ host reads and ∼3.33 × 10^9^ bacterial reads would be required to capture sufficient RNA from both organisms. Thus, to achieve sufficient coverage overall, > 1 × 10^10^ reads would be required for dRNA-Seq of *Chlamydia* and host.

### Infection

Host cells are cultured in Dulbecco’s modified Eagle’s medium (DMEM) (see Materials section) and are infected with *C. trachomatis* at a multiplicity of infection (MOI) of 1 to ensure that 100% of host cells are infected. As cycloheximide (an inhibitor of protein synthesis used to maximize chlamydial yields *in vitro*), is not used throughout this experiment, so the host cells are seeded at ∼60% confluency at the time of infection to ensure continued host cell viability throughout the time-course of the study. Given the time-sensitive nature of the experiment, it is critical to synchronize the initial infection by centrifugation, followed by the removal of dead or non-viable bacterial cells by washing twice with DPBS. HeLa and *Chlamydia* cells are harvested into sucrose phosphate glutamate (SPG) media, and frozen at −80°C prior to RNA extraction.

### RNA Preparation

To ensure high-quality data, dRNA-Seq typically requires a relatively large amount of input RNA. Extreme care must be taken to prevent DNA contamination or RNA degradation, which can be minimized by adhering to the protocol time and temperature requirements, purchasing highly pure and RNase-free reagents, and using RNA-free equipment and consumables. Always conduct RNA work in a clean environment that is partitioned from non-RNA work. Wherever possible we routinely use commercially available kits due to their reliability and reproducibility, however we have carefully optimized the manufacturer’s instructions to suit this protocol.

Total nucleic acid is extracted using a MasterPure™ RNA Purification Kit (Epicenter) according to the manufacturer’s instructions. At this stage it is critical to minimize the delay between host cell lysis and RNA extraction to avoid unwanted degradation. HeLa cells are lysed, host proteins digested, and total nucleic acid precipitated with isopropanol. It is critical to ensure the complete removal of contaminating DNA before proceeding. We have found that two treatments with TURBO DNA-*free*™ DNase (Thermo Fisher) is most effective. We perform three real-time qPCR assays for human targets and one endpoint PCR assay for *C. trachomatis* to confirm DNA removal. The qPCR assays are based on TaqMan^®^ Gene Expression assays (Applied Biosystems) with primer and probe sets targeting beta actin, mitochondrially encoded ATP synthase 6, and eukaryotic 18S rRNA [1]. The endpoint PCR is based on custom-designed primer sets that are specific for *C. trachomatis*, which were designed using PrimerExpress software (Applied Biosystems).

As rRNA constitutes >95% of total RNA, a method of rRNA depletion should be considered to maximize the recovery of mRNA and reduce the sequencing depth required. There are a number of commercial kits available for nuclease digestion and size-selection; this protocol utilizes both a hybridization-based rRNA depletion and poly(A)-depletion step. For hybridization, cDNA oligonucleotides attach to complementary rRNA that is immobilized on magnetic beads; always ensure that the oligonucleotides are compatible with your organism(s) of interest. For this, we combine an equivalent volume of Ribo-Zero beads from both a Human/Mouse/Rat-specific and Gram-negative bacteria-specific Ribo-Zero™ rRNA Removal Kit (Epicenter), enabling simultaneous elimination of both host and bacterial rRNA. It is important to note that this method will not enrich immature mRNAs and non-coding RNAs; specific target enrichment techniques that are outside the scope of this protocol should be considered if these are of experimental interest. Aliquots of rRNA-reduced samples may be then subjected to poly(A) depletion to further enrich host mRNA transcripts and separate mRNA from rRNA contaminants. Poly(A)-depleted and rRNA-depleted eluates can also be further purified before being combined for library construction. The remaining RNA is concentrated and purified with a RNA Clean & Concentrator™-5 kit (Zymo Research). A Bioanalyzer (Agilent) is used to examine the concentration and quality of purified RNA via a capillary electrophoresis-based system.

### Library Preparation and Sequencing

Prior to sequencing, total RNA is converted to cDNA [36]. There are a number of sequencing platforms currently available, including Illumina, SOLID, Ion Torrent, Roche 454, Nanopore, and Pacific Biosciences, and each are suited to specific purposes and should be investigated by the user according to their desired outcome. This stage of the (d)RNA-Seq protocol (library preparation and sequencing) is often outsourced to a commercial enterprise or central sequencing facility so the steps involved are outside the scope of this protocol. Each facility will provide detailed instructions on the quality and quantity of RNA required and the sample preparation guidelines. Nevertheless, here we provide some general guidelines based on our experience.

We generally use the TruSeq Sample Prep Kit for library preparation and sequencing by the Illumina platform. For this, the mRNA is chemically fragmented and primed with random hexamer primers. First-strand cDNA synthesis occurs using reverse transcriptase, followed by second strand cDNA synthesis using DNA polymerase I and RNase H. The cDNA is purified and end-repaired and 3’ adenylated. Adapters containing six nucleotide indexes are ligated to the double-stranded cDNA, which is purified with AMPure XT beads (Beckman Coulter) and enriched via polymerase chain reaction (PCR) amplification. While we suggest that paired-end reads > 50 nucleotides will promote increased fragment randomization and is a good guideline, longer reads will enable greater coverage, reduced multi-mapping, and improved transcript identification [37].

### Data preparation

Sequence data from dRNA-seq comprises cDNA as input from the experiment, with the majority derived from the eukaryotic host (depending on the experimental conditions and system under study). Thus, careful attention is required to accurately segregate the reads from each organism. For paired-end sequencing, host and bacteria read data is generally provided as two FASTQ format files, which are comprised of a unique read identifier, the sequence read, an optional alternate identifier, and the quality scores for each read position. These are examined for possible sample contamination by screening total reads against a sequence database with FastQ Screen (http://www.bioinformatics.babraham.ac.uk/projects/fastq_screen/) (Figure 3). The reads are then checked for quality using FASTQC, a Java-based software that reports several quality control statistics and a judgment on each metric (pass, warn, fail) (Figure 4) [36]. Short reads, low quality reads, and adapters are then removed with Trimmomatic [38]. Other available QC tools available include PRINSEQ [39] and FastX-Toolkit (http://hannonlab.cshl.edu/fastx_toolkit/).

**Figure 3.**
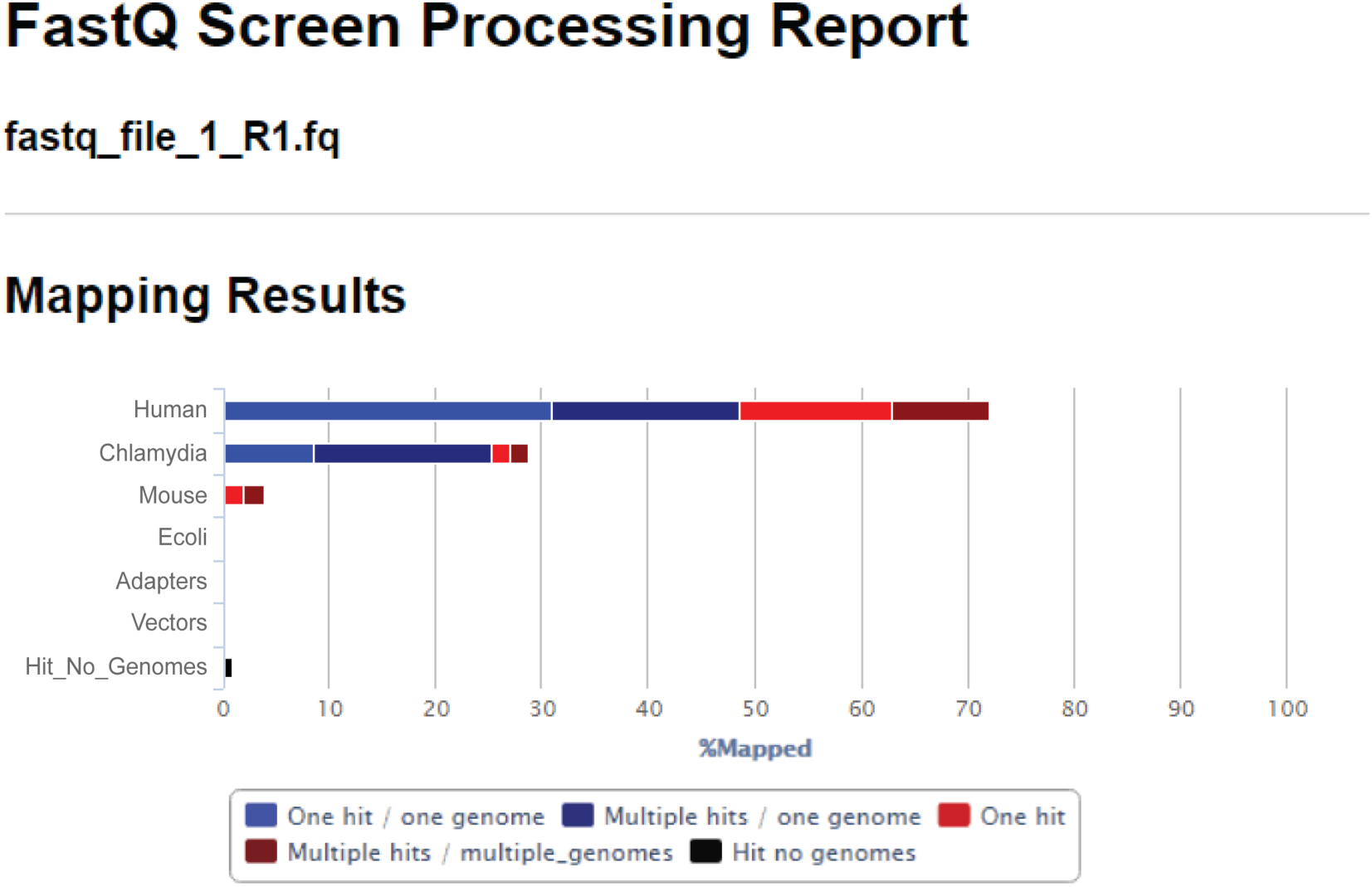
FastQ-Screen processing report of raw host and bacteria FASTQ sequencing reads. As expected, the majority of reads map to the human genome (70%), while 30% of reads map to the *Chlamydia* genome.

**Figure 4.**
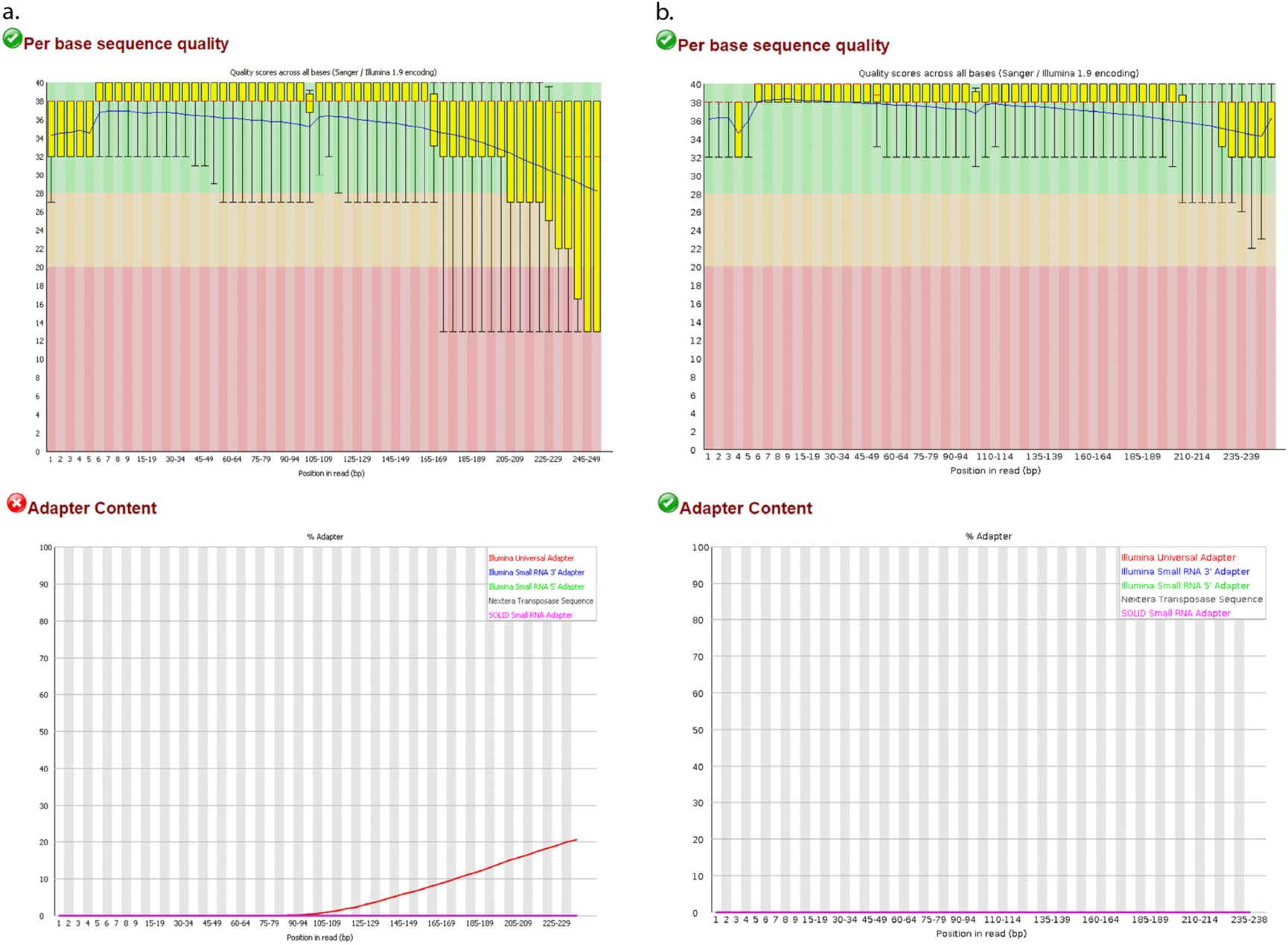
FASTQC report for per base sequence quality and adapter content. **a.** Sequence quality before removal of adapters with TRIMM0MATIC; **b.** Sequence quality after removal of adapters with TRIMM0MATIC.

The organisms of interest and experimental question will dictate which mapping software is most appropriate; we currently use HISAT2 [40], a powerful yet efficient program capable of identifying the splice junctions between exons that are characteristic of eukaryotic data, while the short-read aligner, Bowtie2 [41], is sufficient for bacterial read mapping. Other non-splice-aware aligners for bacterial reads include SEAL [42] and SOAP2 [43], and alternative splice-aware aligners for host reads include MapSplice [44], STAR [45], and Tophat2 [46]. It is important to note that read aligners are an active area of research, with new tools and updates frequently appearing [47].

The combined host and bacteria reads are mapped to the host reference genome with specific settings to preserve unmapped (bacterial) reads, which are then mapped to the bacteria reference genome. Assembly to a reference transcriptome is also possible, but this relies on the accuracy of annotated gene models which may restrict the discovery of novel genes and isoforms; this approach is more suitable when there is no reference genome available. If available for the organisms of interest, reference genomes and the annotation file can be obtained from either NCBI [48], UCSC [49], or Ensembl [50]. Each repository formats these files slightly differently so it is important to obtain both files from the same source. This protocol utilizes the GRCh37 release of the *Homo sapiens* genome and annotation file from Ensembl and the *Chlamydia trachomatis* serovar D genome from NCBI (NC_000117.1).

The resulting alignment files for both host and bacteria are sorted by position (i.e. chromosomal location) with Samtools [51] to produce alignment quality statistics, including the number of mapped reads, the number of mapped first mates and second mates (for reads from paired-end sequencing), reads with multiple hits in the genome, and host reads mapping to exonic, intronic, and intergenic regions. Ideally, > 70% of host reads should map to exonic regions of the genome while less than 5% of reads mapping to intronic regions and less than 1% of reads mapping to intergenic regions [52]. The alignment file is further converted to a BigWig format for the visualization of the number of reads aligned to every single base position in the genome using Integrated Genome Viewer (IGV) (Figure 5) [53], or other visualization tools such as UCSC Genome Browser or JBrowse [54]. Using IGV, the coverage of aligned reads across the genome for both host and bacteria can be visualized to identify genomic regions of high/low coverage that could indicate technical or biological errors, as well as host exon-intron boundaries, splice sites, exon junction read counts, and read strand [55]. The alignment files are then sorted by read name to facilitate feature counting.

**Figure 5.**
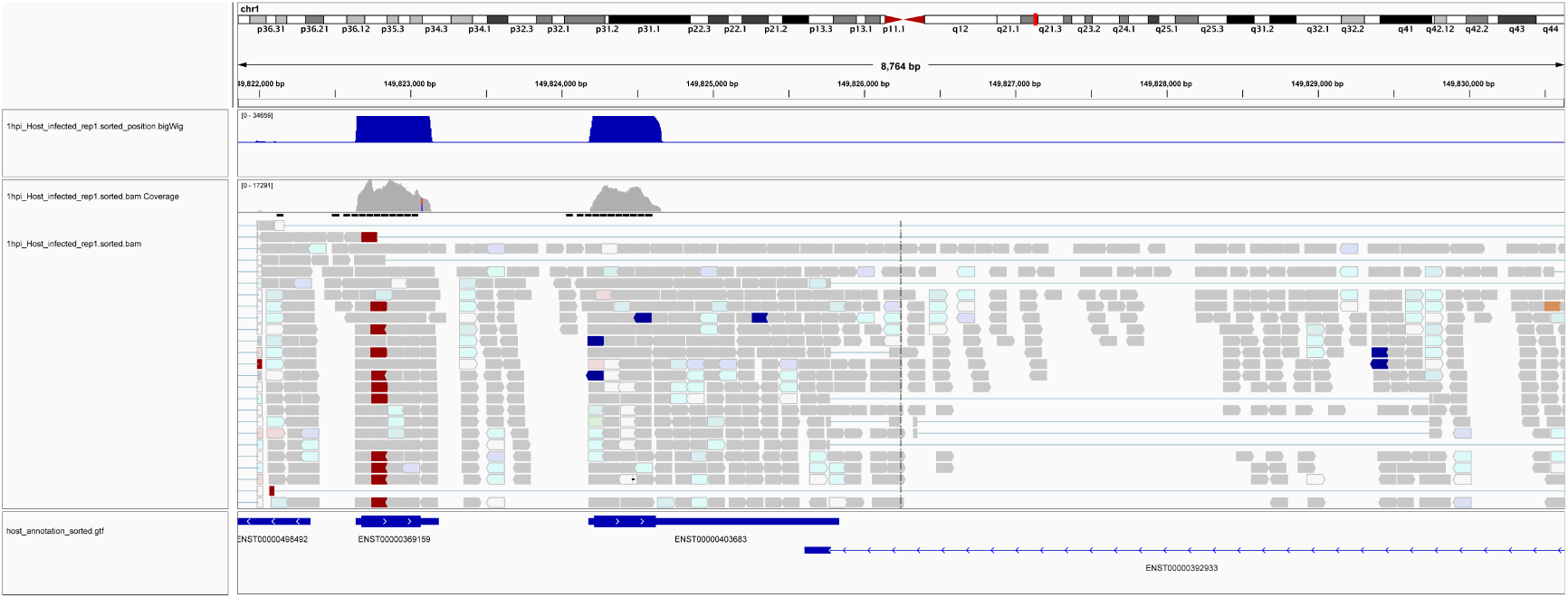
Screen shot of Integrated Genome Viewer (IGV) showing host mapped reads the associated gtf annotation file. The first bar labelled "chr1" indicates which portion of the human genome (or chromosome) is displayed, with the length (8,764 bp) and specific genomic region shown underneath. The graphs shown in blue and gray indicate read coverage and the sequence alignment tracks are shown below this. The bottom row is the gtf annotation file indicating which annotated transcripts the reads are aligning to.

### Feature counting and normalization

Both host and bacteria counts are generated from their respective alignment files using the python wrapper script *htseq-count* from the HTSeq package [56]. This process quantitates the number of reads that align to a biologically meaningful feature such as exons, transcripts, or genes [56], and is guided by the reference annotation file. This protocol describes the quantification of reads on a gene level, where a gene is considered the union of its exons. Any reads that map to several genomic locations are automatically discarded by HTSeq and we generally take a conservative approach to also discard reads that overlap with more than one gene.

Sample read counts are collapsed into a single file containing a matrix of genes (rows) and samples (columns), with one file each for host and bacteria (Figure 6). To minimize statistical noise and enable better adjusted *p-*values, the matrix is pre-filtered so that greater than three counts remain in more than two of the samples [36]. Additionally, the last five lines of the matrix containing statistics for ambiguous counts from *htseq-count* are removed. Raw counts are normalized to minimize technical bias due to transcript length and sequencing depth; there are several normalization methods available, including Reads Per Kilobase Per Million (RPKM) [20], EDASeq [57], Conditional Quartile Normalization (CQN) [58], Upper Quartile (UQ) [59], and Transcripts Per Million (TPM) [60], and each have their benefits. For the host counts we use Trimmed Mean of M-Values (TMM) which corrects for differences in RNA composition and sample outliers, while providing better across-sample comparability [61].

**Figure 6.**
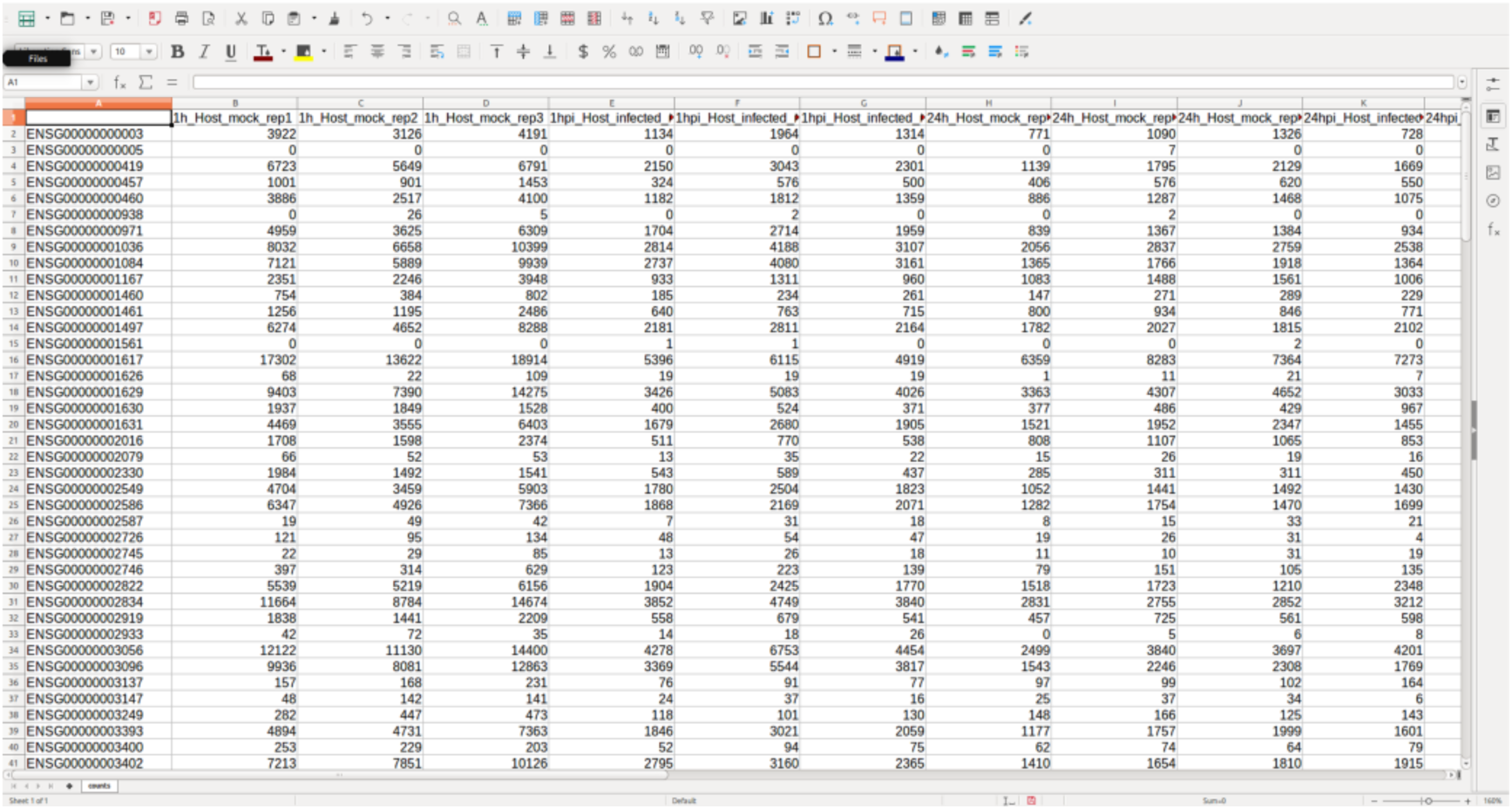
The count matrix. Following read quantification with HTSeq the count files are combined to form the matrix of raw counts for each sample and replicate in the dataset.

### Data analysis

Prior to differential expression analysis, both a Multi-Dimension Scaling (MDS) plot and hierarchical cluster plot is constructed to visualize the distances between samples and help identify problematic and outlying samples (Figures 7 and 8). A metadata table is generated to list the experimental variables, which in turn guides the construction of design and contrast matrices, which are mathematical representations of the experimental design and a description of the relevant treatment comparisons, respectively. This protocol describes a simple design and contrast matrix that allows differential expression comparisons to be made between infected and uninfected cells within each time-point. Additional time-points and other experimental factors would lend themselves to more complex design matrices and may be added by the user if required.

**Figure 7.**
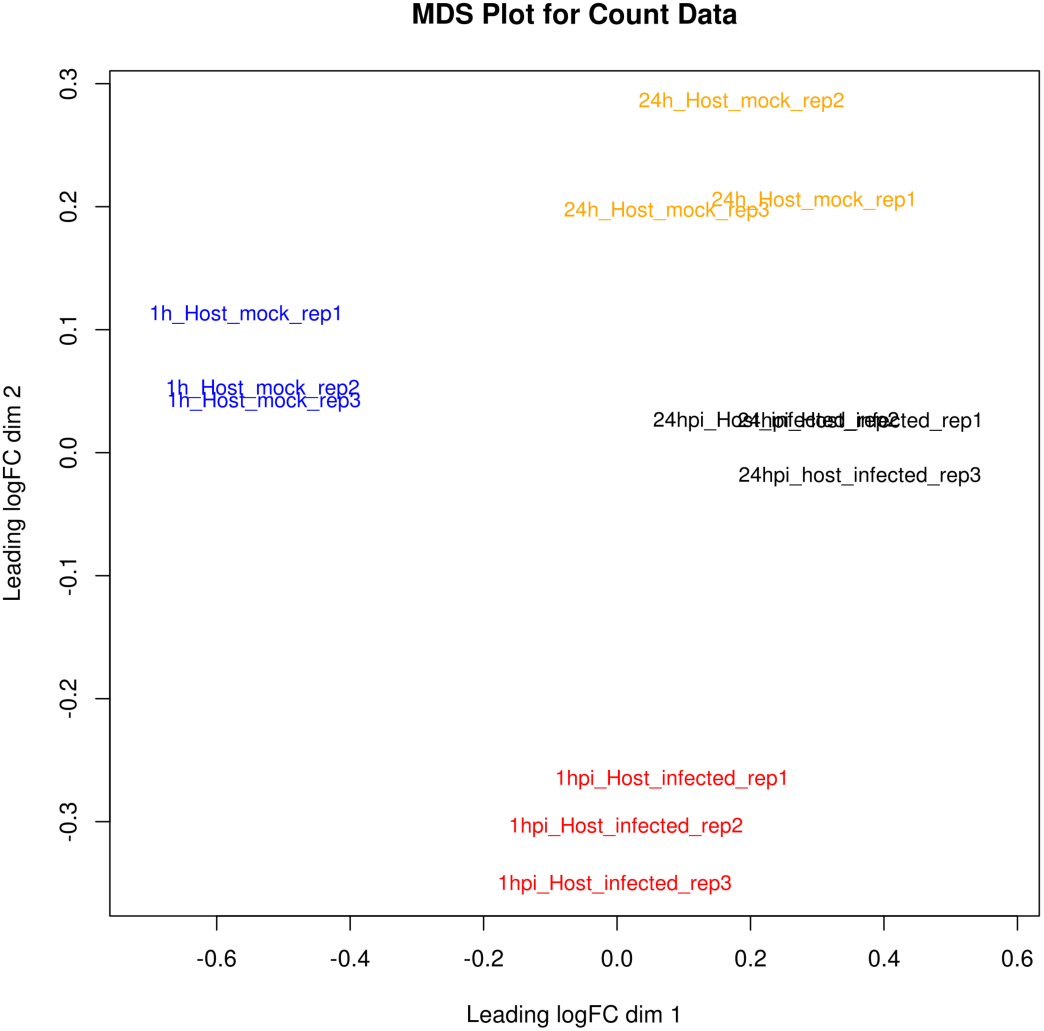
Multi-dimension Scaling Plot (MDS). This is a two-dimensional plot that visualizes the similarity between samples and replicates across conditions. It enables the identification of problematic samples that may obscure the subsequent statistical analysis. In this case, all replicates cluster together as expected.

**Figure 8.**
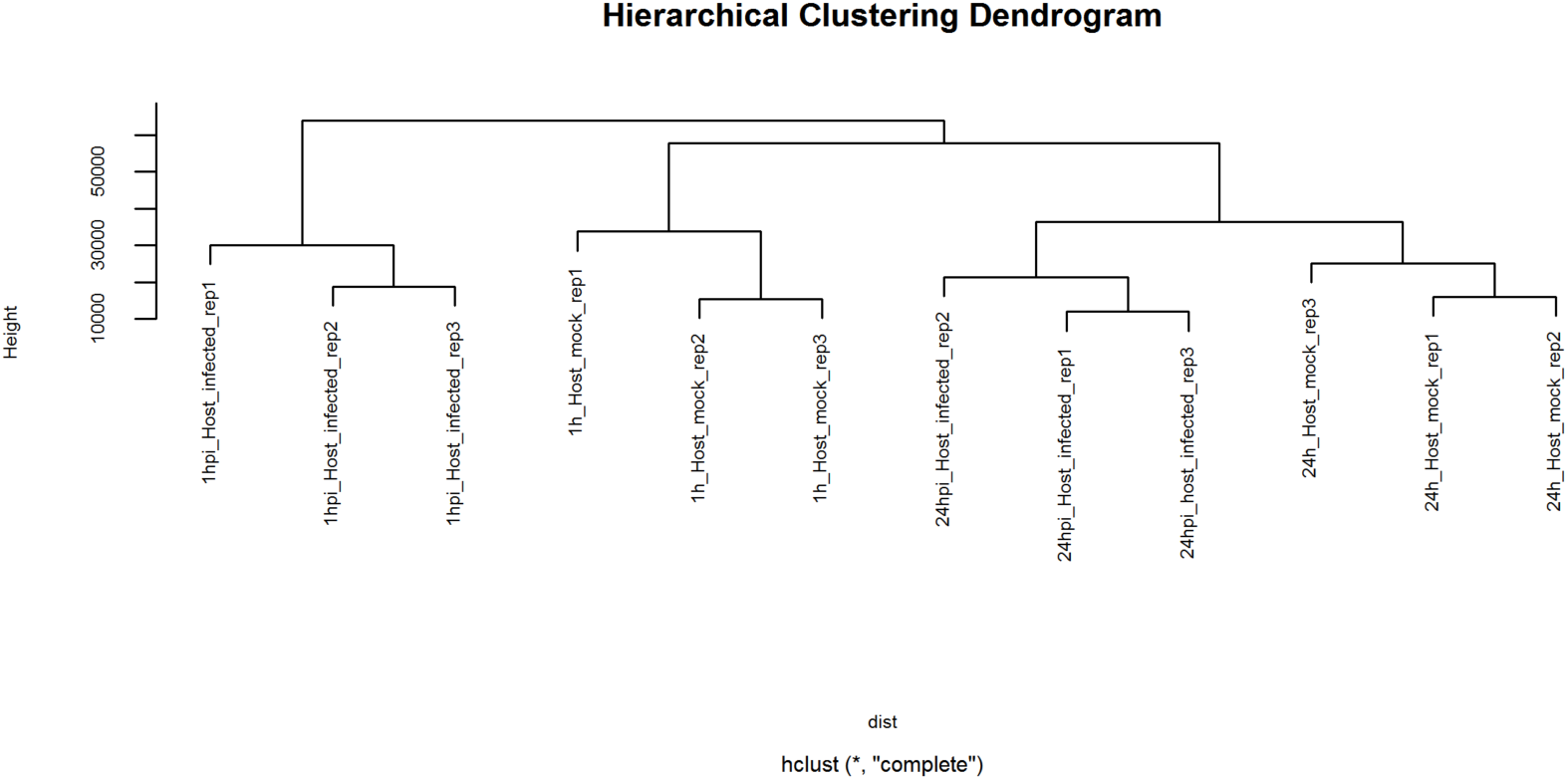
Hierarchical clustering dendrogram. An extension of the MDS plot, the Hierarchical clustering dendrogram illustrates sample similarity. As expected, all replicates for each condition cluster together.

Due to the specific nature of the host and bacteria count data, distinct statistical analyses are required for each. For bacterial transcripts, TPM is the most appropriate measure of relative transcript abundance, but this approach can suffer from biases where the calculated abundance of one transcript can affected other transcripts in the sample. Alternatively, absolute abundance may be calculated with the use of spike-in controls. Whichever method is chosen, these abundances represent a descriptive characterization of the *Chlamydia* transcriptome at the 1- and 24hpi time-points, which is moderately informative on its own, but when analyzed in combination with the interacting host response can allow deeper insight into the host-bacteria interactome.

For the host, differential expression analysis is performed to identify genes that have changed significantly in transcript abundance between infected and time-matched non-infected samples [5]. There are multiple differential expression packages available, each with advantages, including BaySeq [62], Cufflinks [63], DESeq [64], edgeR [65], Salmon [66], and Kallisto [67]. Protocols for the Cufflinks suite and edgeR/DESeq packages have been described previously [5,68]. The majority of these packages model RNA-Seq counts via a negative binomial (NB) distribution and apply various approaches to calculate reliable dispersion estimates. Alternatively, this protocol describes the use of the Limma package which uses linear modeling to describe the expression data for each gene (Limma stands for “Linear Modeling for Microarray Data”) [69]. In contrast to these other RNA-Seq packages, Limma attempts to correctly model the mean-variance relationship between samples to achieve a more probabilistic distribution of the counts (Figure 9). This has proven to be the best method for analyzing both simple and complex experimental designs of dRNA-seq experiments that incorporate different sample types and time-points [70].

**Figure 9.**
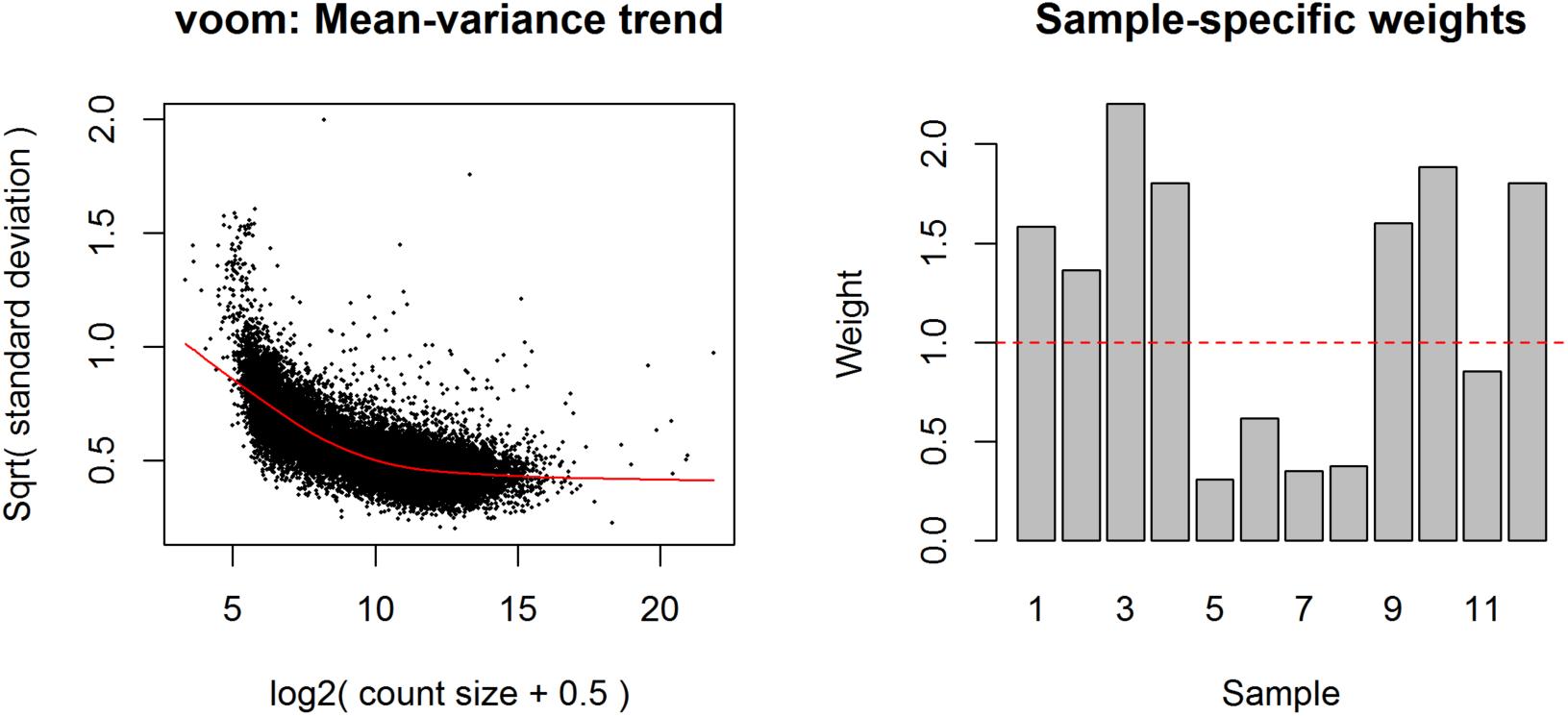
Limma voom plots. The mean variance trend plot displays the gene-wise square-root residual standard deviations plotted against average log-count, with the L0WESS fit represented by the red line. The sample-specific weights are the result of the "voomWithQualityWeights" function and represents the sample-specific quality weights that can be applied to down-weight outlier samples.

The output for the host is a set of differentially expressed genes, applying a False Discovery Rate (FDR) cutoff ≤ 0.05 (i.e. 5% false positives), and at least two-fold up/down-regulation and expression levels greater than 1 percentile in either condition (Table 1). These lists can then be used as input for downstream analysis of the enrichment of gene ontology and metabolic pathways using several tools, including GOSeq [71], DAVID [72], and Ingenuity Pathway Analysis (IPA) Toolkit (QIAGEN Redwood City, www.qiagen.com/ingenuity).

**Table 1.** Statistical output of the differential expression analysis of host reads in R. The first column contains the ENSEMBL ID for the genes, logFC indicates the log fold-change observed, AveExpr is the expression value for each gene, t is the moderated t-statistic, P.Value is the raw *p*-value, adj.P.Val is the false discovery rate-adjusted *p*-value, and B is the log odds that the gene is differentially expressed.

|  | LogFC | AveExpr | t | P.Value | adj.P.Val | B |
| --- | --- | --- | --- | --- | --- | --- |
| ENSG00000003096 | 0.6525066 | 13.791554 | 28.40683 | 1.447731e-10 | 4.921022e-09 | 14.94091 |
| ENSG00000005483 | 0.6818259 | 14.119036 | 28.15577 | 1.574679e-10 | 4.921022e-09 | 14.85045 |
| ENSG00000003436 | 0.6141746 | 15.274943 | 28.06727 | 1.622315e-10 | 4.921022e-09 | 14.60957 |
| ENSG00000004766 | 0.5506978 | 13.234187 | 27.08358 | 2.273831e-10 | 5.172965e-09 | 14.52388 |
| ENSG00000003147 | 4.2364651 | 6.880526 | 22.53363 | 1.289647e-09 | 2.252721e-08 | 11.79033 |
| ENSG00000001630 | -1.8429486 | 11.760177 | -22.19781 | 1.485311e-09 | 2.252721e-08 | 12.61091 |

### Integration of host and bacterial transcriptomic data

The final and most powerful stage of a dRNA-seq experiment is the integration of host and bacterial data to draw biological conclusions that could not have been obtained by investigating each organism in isolation. Following Gene Ontology and pathway analyses, interacting host-bacteria responses can be identified, within and between time-points, to highlight pathogenic mechanisms in the bacterium and the reciprocal regulatory patterns, responses, and transcriptional reprogramming of the host (depending on the system under investigation).

### Application

dRNA-Seq can be used to address a number of experimental questions. Host differential mRNA and miRNA expression, differential exon usage, alternative splicing, and novel transcript and isoform discovery in response to the bacteria can be determined [5], which may be correlated with the transcriptomic response of the bacteria to determine interaction dynamics. These results can be further integrated with other sources of biological input, including genotyping data and epigenetic information (transcription factor binding, histone modification, methylation etc.) to contribute to a systems level definition of host regulatory mechanisms.

## Procedure: Laboratory

### Materials

#### Reagents

HeLa 229 epithelial cells (ATCC^®^ CCL-2.1™) **CRITICAL** All experiments that use human or animal tissues must comply with governmental and institutional guidelines and regulations.

*Chlamydia trachomatis* serovar E **CAUTION** *Chlamydia* is a human bacteria that poses a risk of infection. All work with this organism should be conducted in a class II biosafety cabinet while wearing appropriate personal protective equipment (PPE).

Dulbecco’s modified Eagle’s medium, high glucose, pyruvate (Thermo Fisher, cat. no. 11995-065)

Heat-inactivated fetal bovine serum (Thermo Fisher, cat. no. 10082-147)

Sucrose (Sigma-Aldrich, cat. no. S7903)

Glutamic acid (Sigma-Aldrich, cat. no. 49449)

Di-sodium hydrogen phosphate (Scharlau, cat. no. S003390500)

Sodium di-hydrogen phosphate (Scharlau, cat. no. S003310500)

Dulbecco’s phosphate buffered saline, no calcium, no magnesium (Thermo Fisher, cat. no. 14190-144)

Gentamicin (50 mg/mL) (Thermo Fisher, cat. no. 15750-060) **CAUTION** Harmful if swallowed or inhaled. Causes irritation to the skin. Wear suitable protective clothing when handling.

Streptomycin (Sigma-Aldrich, cat. no. S9137) **CAUTION** Harmful if swallowed. Suspected of damaging fertility or the unborn child. Wear suitable protective clothing when handling.

Ethanol (Sigma-Aldrich, cat. no. E7023) **CAUTION** Highly flammable. Causes skin and serious eye irritation. Handle using appropriate safety equipment.

Isopropanol (Sigma-Aldrich, cat. no. I9516)

TruSeq RNA Sample Prep Kit (Illumina, cat. no. RS-122-2101)

AMPure XT beads (Beckman Coulter, cat. no. A63880)

Poly(A)Purist Mag Purification Kit (Thermo Fisher, cat. no. AM1922)

TURBO DNA-*free*™ kit (Thermo Fisher, cat. no. AM1907)

DEPC-treated water (Thermo Fisher, cat. no. AM9915G)

MasterPure™ RNA Purification Kit (Epicenter, cat. no. MCR85102)

RNA Clean & Concentrator™-5 (Zymo Research, cat. no. R1015)

Ribo-Zero™ rRNA Removal Kit (Human/Mouse/Rat) (Epicenter, cat. no. RZH1046)

Ribo-Zero™ rRNA Removal Kit (Gram-negative bacteria) (Epicenter, cat. no. RZNB1056)

TaqMan^®^ Gene Expression Assay (Thermo Fisher, cat. no. 4453320)

TaqMan^®^ Universal Master Mix (Thermo Fisher, cat. no. 4352042)

#### Equipment

MicroAmp^®^ optical 96-well reaction plates (Thermo Fisher, cat. no. 4306737) MicroAmp^®^ optical adhesive film (Thermo Fisher, cat. no. 4311971)

MicroAmp^®^ optical film compression pad (Thermo Fisher, cat. no. 4312639)

Flask rocker (Grant Instruments, PS-M3D)

DNase and RNase-free 1.5 mL microcentrifuge tubes (Sarstedt, cat. no. 72.692.210) Refrigerated microcentrifuge (Beckman Coulter, 20R)

Centrifuge (Beckman Coulter, X-12R)

Vortex (Scientific Industries, G-560E)

Heating block (x2) (Thermo Fisher, cat. no. 2001Q)

Ice

Ice bucket Timer

Real-time qPCR machine (Applied Biosystems, 7900HT)

Temperature cycler (Bio-Rad, C1000)

Incubator, 37°C, 5% CO_2_ (Sanyo, MCO-19AIC)

Bioanalyzer

6-well plates (Thermo Scientific, cat. no. 140675)

Cell scrapers (Sarstedt, cat. no. 83.1830)

15 mL centrifuge tubes (Thermo Scientific, cat. no. 339652)

#### Reagent Setup

**SPG media** 10 mM sodium phosphate, 250 mM sucrose, 5 mM glutamic acid. Make solution with nuclease-free water at pH 7.4. Store at 4°C for up to six months.

**70% ethanol** Combine 350 mL ethanol with 150 mL nuclease free water. Store at room temperature for up to six months.

**Dulbecco’s modified Eagle’s medium (DMEM)** Media is supplemented with 10% v/v fetal bovine serum, 100 μg/mL streptomycin, 50 μg/mL gentamicin. Store at 4°C for up to six months.

**Streptomycin** Make stock concentration to 100 mg/mL with nuclease-free water. Store as single-use aliquots at −20°C for up to 12 months.

**Sodium phosphate buffer**

**Ribo-Zero™ rRNA Removal Kit** The Ribo-Zero kits are composed of two parts: the magnetic core kit and the rRNA removal reagents. Store the magnetic core kits at 4°C and Mouse/Human/Rat and Gram-negative bacteria rRNA removal reagents at −80°C.

**RNA Clean & Concentrator™-5** Add 48 mL of 100% ethanol to the 12 mL RNA Wash Buffer before use.

### Procedure: Laboratory

#### Seeding and infection

1. Seed 8 × 10^5^ HeLa cells per well in all wells of a six-well plate. Ensure there are three plates per time-point (one infected plate and two non-infected control plates). Incubate plates overnight at 37°C, 5% CO_2_. Following seeding, sit plates on a bench at room temperature for 15 minutes to allow cells to settle, ensuring an even distribution of cells.
2. The next day, infect all plates (except mock-infected control plates) at an MOI of 1.0 to ensure that 100% of the host cells will be infected.
3. Centrifuge plates at 500 x g for 30 minutes at room temperature. Incubate plates at 37°C, 5% CO_2_. Centrifugation is important to synchronize the infections and time-points.
4. Wash cells twice with DPBS and overlay with warm, fresh DMEM media (containing streptomycin, gentamicin, 10% FBS). Incubate at 37°C, 5% CO_2_.

#### Harvesting cells

5 At each time-point, wash cells twice with DPBS and add 1mL DPBS to each well. Harvest cells with a cell scraper and dispense solution into a 15 mL centrifuge tube. Store tubes at −80°C until all time-points are complete. Cells can be stored for up to six weeks at −80°C.

#### Cell lysis

6 Remove centrifuge tubes from −80°C freezer and thaw at room temperature.
7 Pre-set heating block to 65°C.
8 Add 1 μL of 50 μg/mL Proteinase K (MasterPure™ RNA Purification Kit; Epicenter) to 300 μL of Tissue and Cell Lysis Buffer (MasterPure™ RNA Purification Kit; Epicenter) for each sample.
9 Pellet cells by centrifugation at 5,000 x *g* for 30 minutes. Discard the supernatant, leaving ∼25 μL of liquid.
10 Vortex for 10 seconds to resuspend the pellet.
11 Add 300 μL of Tissue and Cell Lysis Solution (containing Proteinase K) to each 25 μL sample and mix thoroughly by vortexing.
12 Incubate tubes in a heating block at 65°C for 15 minutes, vortexing briefly every 5 minutes.
13 Place samples on ice for 3-5 minutes.

#### Total nucleic acid precipitation

14 Add 175 μL of MPC Protein Precipitation Reagent (MasterPure™ RNA Purification Kit; Epicenter) to each 300 μL of lysed sample and vortex vigorously for 10 seconds.
15 Pellet the debris by centrifugation at 10,000 x *g* for 10 minutes at 4°C.
16 Transfer the supernatant (containing total nucleic acid) to a clean 1.5 mL microcentrifuge tube and discard the pellet.
17 Add 500 μL of isopropanol to the recovered supernatant and invert the tube 30-40 times. Do not vortex.
18 Pellet the total nucleic acid by centrifugation at 10,000 x *g* for 10 minutes at 4°C.
19 Carefully pour off the isopropanol without dislodging the pellet.
20 Rinse the pellet twice with 70% ethanol, being careful not to dislodge the pellet. Centrifuge briefly if pellet is dislodged.
21 Remove residual ethanol with a pipette and resuspend the pellet in 35 μL of TE Buffer (MasterPure™ RNA Purification Kit; Epicenter). Samples can be stored in TE Buffer overnight at 4°C.

#### DNA digestion

22 Pre-set heating block to 37°C.
23 Add 4 μL of 10x TURBO Dnase Buffer (TURBO™ DNA-*free*™ Kit; Thermo Fisher) to each 35 μL sample.
24 Add 1 μL of TURBO™ DNase to each sample. Gently flick the tubes to mix and pulse-spin to distribute liquid to the bottom of the tube. Increase DNase volume to 2-3 μL if digesting a higher amount of DNA. Alternatively, add half the DNase to each reaction, incubate for 30 minutes, then add the remainder of the enzyme and incubate for another 30 minutes.
25 Incubate tubes in a heating block at 37°C for 30 minutes.
26 After incubation, add 8 μL of DNase Inactivation Solution (TURBO™ DNA-*free*™ Kit; Thermo Fisher) and incubate tubes at room temperature for 5 minutes, mixing occasionally. Environments colder than 22°C can reduce the inactivation of TURBO™ DNase. Move tubes to a heating block to control the temperature if necessary.
27 Centrifuge tubes at 10,000 x *g* for 1.5 minutes.
28 Transfer supernatant (containing the RNA) to a fresh 1.5 mL microcentrifuge tube.

#### Validation of DNA removal by RT qPCR

29 Remove 5 μL from each RNA sample and place in a clean 1.5 mL microcentrifuge tube.
30 Add 95 μL of nuclease-free water (1:20 dilution).
31 Prepare enough RT qPCR master mix to assay each sample in triplicate, as well as non-template controls:
20x TaqManâ Gene Expression Assay: 1 μL
2x TaqManâ Gene Expression Master Mix: 10 μL
RNase-free water: 5 μL
32 Cap the tube and invert the tube several times to mix the reaction components. Pulse vortex.
33 Aliquot 15 μL of master mix into individual wells of a 96-well reaction plate (Thermo Fisher). Include a triplicate reaction for each sample and non-template controls.
34 Add 5 μL of each diluted template to appropriate wells and gently tap plate on benchtop to distribute contents to the base of the well.
35 Place adhesive film (Thermo Fisher) over the plate and seal with compression pad (Thermo Fisher). If any bubbles are visible in the wells or liquid is present on the sides of the wells, centrifuge plate at 500 x *g* for 2 minutes. Do not touch the film with bare hands at any point.
36 Place plate in RT-PCR machine and run assay according to the following cycling conditions: Hold: 95°C, 10 minutes Cycle (40x): 95°C, 15 seconds 60°C, 1 minute.

#### rRNA depletion

37 Pre-set one heating block to 68°C and one at 50°C.
38 Remove Ribo-Zero™ rRNA Removal Magnetic Core Kit from 4°C and allow to warm to room temperature. Remove Human/Mouse/Rat and Gram-negative bacteria components of the Ribo-Zero™ rRNA Removal kits from −80°C and thaw on ice. Do not place the Ribo-Zero™ Magnetic Core Kit on ice.
39 Vigorously mix magnetic beads (Ribo-Zero™ rRNA Removal kit; Illumina) for 20 seconds by vortexing.
40 Carefully pipette 65 μL of magnetic beads into 2 mL microsphere wash tubes (Ribo-Zero™ rRNA Removal kit; Illumina); two tubes per sample. Store unused beads at 4°C. Do not place the magnetic beads on ice.
41 Centrifuge microspheres at 12,000 x *g* for 3 minutes. Carefully remove supernatant without dislodging the pellet. The supernatant contains sodium azide.
42 Wash the microsphere wash tubes by adding 130 μL of microsphere wash solution (Ribo-Zero™ rRNA Removal kit; Illumina) to each tube. Vortex vigorously.
43 Centrifuge microsphere wash tubes at 12,000 x *g* for 3 minutes. Carefully remove supernatant without dislodging the pellet.
44 Add 65 μL of microsphere resuspension solution (Ribo-Zero™ rRNA Removal kit; Illumina) to each tube and vortex vigorously until a homogenous suspension is produced.
45 Add 1 μL of RiboGuard RNase inhibitor (Ribo-Zero™ rRNA Removal kit; Illumina) to each tube. Mix by vortexing for 10 seconds and set aside (at room temperature). Avoid creating air bubbles when adding RNase inhibitor.
46 Treat two aliquots of each sample with Ribo-Zero rRNA removal solution (Ribo-Zero™ rRNA Removal kit; Illumina) according to the following preparation (two removal preps per sample): RNase-free water (Ribo-Zero™ rRNA Removal Kit): 1 μL Ribo-Zero Reaction Buffer: 4 μL RNA sample: 25 μL Ribo-Zero rRNA Removal Solution (Gram negative bacteria kit): 5 μL Ribo-Zero rRNA Removal Solution (Human/Mouse/Rat kit): 5 μL Fully mix the samples by pipette-mixing 10-15 times.
47 Gently mix the reactions and incubate at 68°C for 10 minutes in heating block. Return the Ribo-Zero reaction buffer to −80°C.
48 Remove the microsphere wash tubes from the heating block and incubate at room temperature for 15 minutes.
49 Vortex the microsphere wash tubes at medium speed for 20 seconds to ensure a homogenous slurry.
50 Pipette hybridized RNA sample to the resuspended microsphere wash tubes, pipette-mixing 10-15 times to mix. Immediately vortex the microsphere wash tubes at medium speed for five seconds. The washed magnetic beads must be at room temperature for use in this step. The order of the addition (hybridized RNA *to* the magnetic beads) is critical for rRNA removal efficiency.
51 Incubate microsphere wash tubes at room temperature for 10 minutes. Vortex at medium speed for 5 seconds, every 3-4 minutes.
52 Following incubation, mix samples again by vortexing at medium speed for 5 seconds.
53 Incubate samples in heating block at 50°C for 10 minutes.
54 Transfer the RNA-microsphere suspension to a Microsphere Removal Unit (Ribo-Zero™ rRNA Removal kit; Illumina) and centrifuge at 12,000 x *g* for 1 minute at room temperature. Save the eluate and discard the removal unit. At this stage, the eluate should be ∼100 μL.

#### Purification of rRNA-depleted samples

55 Add 2 volumes of RNA Binding Buffer (RNA Clean & Concentrator™-5; Zymo Research) to each volume of RNA sample and mix well. The minimum recommended sample volume for use with this kit is 50 μL.
56 Add 1 volume of 100% ethanol to the mixture and mix well.
57 Transfer the mixture to a Zymo-Spin IC column (RNA Clean & Concentrator™-5; Zymo Research) in a collection tube and centrifuge at 12,000 x *g* for 1 minute. Discard the flow-through.
58 Combine two reactions of the same sample to one column and spin multiple times until the entire mixture passes through the column. The column capacity is 5 μg of RNA.
59 Add 400 μL of RNA Prep Buffer (RNA Clean & Concentrator™-5; Zymo Research) to the column and centrifuge at 12,000 x *g* for 1 minute. Discard the flow-through.
60 Add 800 μL of RNA Wash Buffer (RNA Clean & Concentrator™-5; Zymo Research) to the column and centrifuge at 12,000 x *g* for 1 minute. Discard the flow-through.
61 Add 400 μL of RNA wash buffer to the column and centrifuge at 12,000 x *g* for 1 minute. Discard the flow-through.
62 Centrifuge the column in an emptied collection tube at 12,000 x *g* for 2 minutes. Carefully remove the column from the collection tube and transfer to a new RNase-free microcentrifuge tube.
63 Add 20 μL of DNase/RNase-free water directly to one column matrix and let stand for 1 minute at room temperature. Centrifuge at 10,000 x *g* for 30 seconds. The eluted RNA can be used immediately or stored at −80°C.

#### Library Preparation and Sequencing

The method of library preparation is dependent on the sequencing platform used and thus is outside the scope of this protocol. Commercial sequencing enterprises and sequencing centers will provide detailed guidelines on the library preparation and sample submission guidelines that begin with the isolation of pure mRNA as described above. Following sequencing, the user will be provided with a series of raw FASTQ sequence files that serve as the input for the following bioinformatics section.

## Procedure: Bioinformatics

### Materials

#### Hardware requirements

The analysis of dRNA-seq experiments is a computationally intensive process that requires the manipulation of gigabytes of data. Access to a computer cluster, core facility, or cloud service is recommended to expedite the analysis and free up resources on the local system.

#### Operating system

This protocol provides commands that are designed to run on a Unix-based operating system such as Linux or Mac OS. The protocol was specifically designed to run on the Ubuntu 16.04.1 operating system on a Linux machine. Please ensure you have administrative rights.

#### Command line nomenclature

This protocol assumes a basic understanding of both the Linux command line interface and the R statistical computing environment. All Linux commands are shown following a dollar sign ($), while R commands are shown following a greater-than sign (>):

$ linux command

> R command

#### Software requirements

- FastQ Screen: Contamination screening (http://www.bioinformatics.babraham.ac.uk/projects/fastqscreen/)
- FASTQC: Sequence quality control tool (http://www.bioinformatics.babraham.ac.uk/projects/fastqc/)
- Trimmomatic: FASTQ sequence file trimming (http://www.usadellab.org/cms/?page=trimmomatic)
- HISAT2: Graph-based alignment of sequences to genomes (https://ccb.jhu.edu/software/hisat2/index.shtml)
- Samtools: Manipulation of sequence alignments and mapped reads (http://www.htslib.org)
- Bedtools genomecov and bedGraphToBigWig: genomic analysis tools (http://bedtools.readthedocs.io/en/latest/)
- Integrated Genome Viewer (IGV): Alignment and visualization tool (https://www.broadinstitute.org/ig)
- HTSeq: read counting (http://www-huber.embl.de/users/anders/HTSeq/doc/overview.html)
- R statistical computing environment (https://www.r-project.org)
- Bioconductor packages: edgeR, limma, org.Hs.eg.db, GenomicFeatures, and their dependencies (see below)
- Bowtie2: Short-read aligner (http://bowtie-bio.sourceforge.net/index.shtml).

Always check that you are downloading and installing the latest version of each piece of software and consult the official user guide for more in-depth guidelines and options for troubleshooting any errors that may arise.

#### Samples and filenames

The protocol is arranged so that identical naming conventions are used for each sample and condition. For example, “1hpi_Host_infected_rep1” indicates that the sample relates to the first replicate of host cells infected with *Chlamydia* at the 1 hpi time point and “1hpi_Host_uninfected_rep1” indicates the first replicate of host cells only (i.e. uninfected host cells) at the 1 hpi time point. Conversely, the samples relating to the bacteria are named, “1hpi_Bacteria_rep1”. While subsequent replicates for both host and bacteria would have names ending in “rep2” and “rep3”, for conciseness this protocol describes the commands using “1hpi_Host_infected_rep1” as an example and it is expected that the user will repeat the process for the remaining replicates and samples. The filenames associated with raw FASTQ sequence files will depend on the sequencing facility pipeline and in this protocol are named “fastq_file_1_R1.fq”, where “R1” indicates read number one of paired end reads (the corresponding read file would be “fastq_file_1_R2.fq”. In some cases an output directory is required, which is noted as “<output_directory>” for the user to input their working directory of choice (without the “<>” symbols). Finally, reference, annotation, and gene info files are prefixed with the relating organism, i.e. “host_reference.fa” indicates a FASTA file containing the host reference genome.

##### Equipment setup

Download and install the following software. Check the developer website to ensure you are installing the latest version and for further information about dependencies and prerequisites.

#### Create a directory to install program executables and add to PATH

$ mkdir $HOME/bin

$ export PATH=$HOME/bin: $PATH

$ echo "export PATH=$HOME/bin:$PATH" >> ∼/.bashrc

#### FastQ-Screen installation

$ wget

http://www.bioinformatics.babraham.ac.uk/projects/fastqscreen/fastqscreenv0.9.3.tar.gz

$ tar -zxf fastq_screen_v0.9.3.tar.gz

$ cd fastq_screen_v0.9.3

$ cp fastq_screen $HOME/bin

#### FastQC installation

$ sudo apt-get install default-jre

$ wget http://www.bioinformatics.babraham.ac.uk/projects/fastqc/fastqc_v0.11.5.zip

$ unzip fastqc_v0.11.5.zip

$ cd FastQC/

$ chmod 755 fastqc

$ cp fastqc $H0ME/bin

**Trimmomatic installation**

$ wget http://www.usadellab.org/cms/uploads/supplementary/Trimmomatic/Trimmomatic-0.36.zip

$ unzip Trimmomatic-0.36.zip

$ cd Trimmomatic-0.36

$ cp trimmomatic $HOME/bin

#### HISAT2 installation

$ wget ftp://ftp.ccb.jhu.edu/pub/infphilo/hisat2/downloads/hisat2-2.0.5-Linux_x86_64.zip

$ unzip hisat2-2.0.5-Linux_x86_64.zip

$ cd hisat2-2.0.5

$ cp hisat2 $H0ME/bin

$ cp hisat2-build $H0ME/bin

#### Samtools installation

$ sudo apt-get install samtools

#### Bedtools installation

$ sudo apt-get install bedtools

#### bedGraphToBigWig installation

$ mkdir bedGraphToBigWig

$ cd bedGraphToBigWig

$ wget −0 bedGraphToBigWig https://github.com/ENC0DE-DCC/kentUtils/blob/v302.1.0/bin/linux.x86_64/bedGraphToBigWigraw=true$chmod755bedGraphToBigWig$cpbedGraphToBigWig$H0ME/bin

#### IGV and IGVtools installation

$ wget http://data.broadinstitute.org/igv/projects/downloads/IGV_2.3.88.zip

$ unzip IGV_2.3.88.zip

$ wget http://data.broadinstitute.org/igv/projects/downloads/igvtools2.3.88.zip

$ unzip igvtools_2.3.88.zip

$ cd igvtools_2.3.88

$ cp igvtools $H0ME/bin

#### HTSeq installation

$ sudo apt-get install build-essential python2.7-dev python-numpy python-matplotlib

$ wget --no-check-certificate https://pypi.python.org/packages/source/H/HTSeq/HTSeq-0.6.1p1.tar.gz

$ tar -zxvf HTSeq-0.6.1p1.tar.gz

$ cd HTSeq-0.6.1p1

$ python setup.py build

$ sudo python setup.py install

$ cd scripts

$ cp htseq-count $H0ME/bin

#### R and Bioconductor package installation

$ sudo apt-get install libcurl4-openssl-dev libxml2-dev

$ sudo apt-get update

$ echo “deb https://cran.rstudio.com/bin/linux/ubuntuxenial/” |sudo tee -a /etc/apt/sources.list

$ sudo apt-key adv --keyserver keyserver.ubuntu.com --recv-keys E084DAB9

$ sudo add-apt-repository ppa:marutter/rdev

$ sudo apt-get update

$ sudo apt-get upgrade

$ sudo apt-get install r-base

Open R and install Bioconductor packages using the BiocLite installation tool. All packages dependencies will automatically be installed.

$ R

>. source(“http://bioconductor.org/biocLite.R”)

> biocLite(“BiocUpgrade”)

> biocLite(c(“org.Hs.eg.db”, “edgeR”, “limma”, “GenomicFeatures”))

#### Bowtie2 installation

$ wget https://sourceforge.net/projects/bowtie-bio/files/bowtie2/2.2.9/bowtie2-2.2.9-linux-x86_64.zip

$ unzip bowtie2-2.2.9-linux- x86_64.zip

$ cd bowtie2-2.2.9-linux-x86_64

$ cp bowtie2 $HOME/bin

$ cp bowtie2-build $HOME/bin

*File preparation*

#### Download reference genomes and gene model annotation files

For the eukaryotic host, download the *Homo sapiens* reference genome and GTF annotation file from Ensembl: http://asia.ensembl.org/info/data/ftp/index.html and rename files to “host_reference.fa” and “host_annotation.gtf”, respectively. Download the *Homo sapiens* gene info file from: ftp://ftp.ncbi.nlm.nih.gov/gene/DATA/GENEINFO/Mammalia/Homosapiens.geneinfo.gz and rename file as “host_gene.info”. For *C. trachomatis,* download the reference genome in FASTA format from NCBI: http://www.ncbi.nlm.nih.gov/nuccore/NC_000117 and rename the file to “bacteria_reference.fa”. Download the *C. trachomatis* GTF (000590675) file from bacteria.ensembl.org/info/website/ftp/index.html and rename file to “bacteria_annotation.gtf”. Save all files to your working directory. Reference genomes and annotation files should be obtained from the same repository to ensure consistent formatting and nomenclature.

### Method: Host

#### 1. Examine a subset of FASTQ sequence files for contamination using FastQ Screen

$ fastq_screen -aligner bowtie2 fastq_file_1_R1 .fq

This command will run FastQ Screen on the chosen FASTQ file, checking against locally pre-built databases for possible sources of contamination. “fastq_screen” runs the software, --aligner Bowtie2 specifies the aligner used to create the databases, and “fastq_file_1_R1.fq” is the input file. This step should be repeated for a random number of samples. To generate a database, the genomes of each species which to test against should be downloaded. Using the host and bacterial genomes already downloaded from the earlier steps, Bowtie2 (or other aligners) are used to build an index (See section 4 below). Once built, the location and index should be added to the FastQ Screen configuration file. See Figure 3.

#### 2. Check the quality of FASTQ sequences using FASTQC

$ fastqc -noextract -o <output_directory> fastq_file_1_R1.fq

In this command, “-noextract” tells FASTQC to not uncompress the output file, while “-o” defines the output directory. “fastq_file_1_R1.fq” is the FASTQ sequencing file. These commands produce a quality report with results saved to the directory defined by “<output_directory>”. The results are reported in both illustrated form (the “fastqc_report.html” file) and text form (the “summary.txt” file). Repeat for all FASTQ files. As the FASTQ files are derived from total RNA sequencing, this step includes both host and bacterial sequences. See Figure 4.

#### 3. Remove sequencing adapters and low quality reads using Trimmomatic

$ java-jar trimmomatic-0.36.jar PE -threads 6 -phred33 fastq_file_1_R1.fq fastq_file_1_R2.fq

fastq_file_1_R1_paired_trimmed.fq fastq_file_1_R1_unpaired_trimmed.fq fastq_file_1_R2_paired_trimmed.fq fastq_file_1_R2_unpaired_trimmed.fq ILLUMINACLIP:adapters.fa:2:30:10 LEADING:3 TRAILING:3 SLIDINGWIND0W:4:15 MINLEN:36

Run this command from the Trimmomatic installation directory. The command specifes PE as paired-end data, 6 threads and the FASTQ files are encoded with Phred + 33 quality scores. “fastq_file_1_R1.fastq.gz” and “fastq_file_1_R2.fastq.gz” specify the input FASTQ files to use. As paired-end data is inputted, four output files are needed to store the reads. Two ‘paired’ files from which both reads survived after processing, and two ‘unpaired’ files from which a single read survived, but the corresponding mate did not. “ILLUMINACLIP:adapters.fa” uses the “adapters.fa” file containing sequences and names of commonly used adapters to remove. “2:30:10” are three parameters used in the ‘palindrome’ mode of Trimmomatic to identify the supplied adapters, regardless of their location within a read. For a detailed description of the best use of these three parameters, consult the Trimmomatic manual. “LEADING:3” and “TRAILING:3” removes a base from either the start or end position if the quality is below “3”. “SLIDINGWIND0W:4:15” performs trimming based on a sliding window method. “4” is the window size and “15” is the required average quality. By examining multiple bases, if a single low quality base is encountered, it will not cause high quality data later in the read to be removed. Finally, “MINLEN:36” removes any remaining reads that are less than 36 bases long. Repeat for all FASTQ files. As above, this step includes both host and bacterial sequences.

#### 4. Build host transcriptome index and align host sequence reads to reference using HISAT2

$ hisat2-build host_reference.fa host_reference.index

$ hisat2 -x host_reference.index -un-conc pair1_unmapped.fastq -1 fastq_file_1_.R1.trim.fq -2

fastq_file_1_.R2.trim.fq | samtools view -bS - > accepted_hits.bam

This “-x” specifies the path to the index previously built by Bowtie2: “host_reference.index”. The “— un-conc” argument tells HISAT2 to write a fastq rile containing all unmapped reads (“pair1_unmapped.fastq”), and “-1” and “-2” specify the paired-end fastq file mates. The “samtools view -bS -” argument converts to the output file from SAM to BAM format. Ensure that the HISAT2 output files, “accepted_hits.bam” and “pair1_unmapped.fastq” are preserved in the working directory as these are required for bacterial read mapping.

#### 5. Sort BAM files generated by HISAT2 by both name and position using Samtools

$ samtools sort accepted_hits.bam -o 1hpi_Host_infected_rep1.sorted_position

$ samtools sort -n accepted_hits.bam -o 1hpi_Host_infected_rep1.sorted_name

The first command takes the “accepted_hits.bam” file and sorts it by position, with the output file called “1hpi_Host_infected_rep1.sorted_position”. In the second command, “-n” tells Samtools to sort the “accepted_hits.bam” file by name, with the output file called “1hpi_Host_infected_rep1.sorted_name”. Repeat for all BAM files.

#### 6. Convert ‘sorted by position’ BAM file to BigWig format

$ samtools faidx host_reference.fa

$ cut —f1,2 host_reference.fa.fai > host_reference.genome

$ bedtools genomecov —split —bg —ibam 1hpi_Host_infected_rep1.sorted_position.bam —g

host_reference. genome > 1hpi_Host_infected_rep 1 .sorted_position.bedGraph $ bedGraphT oBigWig 1hpi_Host_infected_rep1 .sorted_position.bedGraphhost_reference. genome 1hpi_Host_infected_rep1 .sorted_position.bigWig

The first command indexes the reference genome by creating a “host_reference.fa.fai” output file. The second command extracts the first two fields (sequence ID and sequence length) to generate the “host_reference.genome” file. The third command generates a histogram illustrating alignment coverage according to the reference genome. The “-split” argument tells *genomecov* to take into account spliced BAM alignments (as we used the splice-aware aligner HISAT2 for the host reads), while the “-bg” argument tells *genomecov* to report genome-wide coverage in bedGraph format. “ibam 1hpi_Host_infected_rep1.sorted_position.bam” is the input file in BAM format, “-g host_reference.genome” is the reference genome in FASTA format, and “1hpi_Host_infected_rep1.sorted_position.bedGraph” is the output file in bedGraph format. The fourth command converts this bedGraph file to BigWig format for use with IGV (below).

#### 7. Index the ‘sorted by position’ BAM file for visualization in IGV

$ samtools index 1hpi_Host_infected_rep1.sorted_position.bam

This command takes the “1hpi_Host_infected_rep1.sorted_position” bam file created above and creates an indexed “1hpi_Host_infected_rep1.sorted_position.bai” file for use with IGV.

#### 8. Index the GTF file for visualization in IGV

$ igvtools sort host_annotation.gtf host_annotation_sorted.gtf

The first command sorts the GTF file, specifying an input file and output file. The second command creates an index of the sorted GTF file.

#### 9. Visualize alignments with IGV

$ java -jar igv.jar

Run this command from the IGV installation directory. Within the software, load the “host_reference.fa” reference genome by clicking on *Genomes* and then *Load genome from file.* Load the sample BAM files from step 9, the indexed GTF annotation file (step 10) and the bigwig file (step 8) by clicking on *File,* then *Open.* Inspect the mapped reads and visualize their alignment to the reference genome to ensure whole genome coverage and that they align with exons as defined by the GTF file. The sample BAM files and index files must be in the working directory. See Figure 5.

#### 10. Create count matrix with HTSeq

$ htseq-count -s no -a 10 -r name -f bam 1hpi_Host_infected_rep1.sorted_name.bam

Host_annotation.gtf > 1hpi_Host_infected_rep1.sorted_name.count

This command calls the *htseq-count* python wrapper script which performs the gene-level counts. “-s no” indicates that the reads are unstranded and “-a 10” sets the minimum mapping quality for a read to be counted as 10. “-r name” indicates that the input file is sorted by name and “-f bam” indicates that this input file is in BAM format and named 1hpi_Host_infected_rep1.sorted_name.bam”. “host_annotation.gtf” is the GTF annotation file from Ensembl and “1hpi_Host_infected_rep1.sorted_name.count” is the output file produced. Repeat command for each BAM file, which will produce a series of text files counting the gene-level reads for each sample. The last five lines of each file contain a list of reads that were not counting due to alignment ambiguities, multi-mapping, or low alignment quality.

#### 11. Set working directory in R

> R

> data <- setwd(…)

The first command opens R, and the second command sets the working directory to the folder location containing all the relevant files from step 12. Replace “…” with the complete path to this location, for example: setwd(“/home/username/dRNA-Seq/HTSeq_counts”).

#### 12. Create data frame in R containing experiment metadata

> group <- factor(c(rep(“ 1hpi_mock”, 3), rep(“ 1hpi_infected”, 3), rep(“24hpi_mock”, 3), rep(“24hpi_infected”, 3)))

This creates the metadata table containing all the experimental variables, including sample name, treatment, and time point.

#### 13. Combine count files into a DGEList in R

> library(edgeR)

> counts.host <- readDGE(list.files(pattern = “.count”), data, columns = c(1,2))

A DGEList is an R object from the edgeR package that efficiently compiles the count dataset and experimental variables that is fed into subsequent downstream analyses. The first command loads edgeR into the current R workspace. The second command creates a variable called counts_host, which is a DGEList containing all data from columns 1 and 2 from all files in the current working directory ending in “.count”. See Figure 6.

#### 14. Remove the last five rows from the count matrix

> counts.host$counts <- counts.host$counts[1:(nrow(counts.host$counts)-5),]

This command removes the last five rows of the count matrix, which contain a summary of the ambiguous and non-counted reads from htseq-count.

#### 15. Filter counts to exclude low expressing genes

> counts.host$counts <- counts.host$counts[rowSums(counts.host$counts > 3) > 2,]

This command returns the count matrix so that there are greater than three reads in at least two replicates across all of the samples. Any non-conforming samples are removed.

#### 16. Inspect the count matrix

> head(counts.host$counts, 20)

> dim(counts.host$counts)

It is often helpful to visualize the count matrix at this point to confirm that it is formatted correctly and that there are no errors. The second command provides the matrix dimensions, which is useful for determining the number of genes remaining following independent filtering.

#### 17. Apply TMM normalization to the raw counts

> counts.host <- calcNormFactors(counts.host)

TMM normalization factors are calculated and incorporated into the DGEList object.

#### 18. Create the design matrix and define the contrasts of interest

> design <- model.matrix(∼0 + group)

> rownames(design) <- colnames(counts.host$counts)

> contrasts <- makeContrasts(“Host_1hpi” = group1hpi_infected -group1hpi_mock,

“Host_24hpi” = group24hpi_infected - group24hpi_mock, levels= design)

These commands define the design matrix and the contrasts of interest to enable differential expression to be calculated. In this case, the contrasts we are interested are the host genes differentially expressed when infected versus the host genes differentially expressed when uninfected at the 1 hpi time point and at the 24 hpi time point. More complex experimental designs that include multiple samples, time points, batch effects, and treatments are possible and are explained in detail in the Limma User’s Guide [73].

#### 19. Apply voom transformation to normalized counts

> library(limma)

> png(“host_voom.png”)

> y <- voomWithQualityWeights(counts = counts.host, design = design, plot = TRUE)

> dev.off()

This command applies a voom transformation to the counts, by converting them to log-counts per million with associated precision weights [70]. We generally extend this by using the *voomWithQualityWeights* function, which applies sample-specific weights to down-weight any outlier samples. This can be especially useful if outliers were identified in the MDS plot constructed in step 21 (below). This function takes as input the normalized count matrix (“counts_host”) and design matrix (“design”) and outputs two quality control plots: an estimation of the mean variance relationship and the sample-specific weights that were applied. The output figure “host_voom.png” is generated containing the voom transformation plots. See Figure 9.

#### 20. Construct an MDS plot to identify any outlier samples

> plot.colors <- c(rep(“blue”, 3), rep(“red”, 3), rep(“orange”, 3), rep(“black”, 3))

> png(“host_MDS.png”)

> plotMDS(counts.host, main = “MDS Plot for Count Data”, labels = colnames(counts.host$counts), col = plot.colors, cex = 0.9, xlim=c(-2,5))

> dev.off()

These commands generate a Multi-Dimensional Scaling (MDS) plot by taking the host DGEList as input (“counts_host”). The MDS plot allows the visual inspection of sample proximities to highlight possible batch effects and sample outliers that may need to be addressed. The MDS plot is saved to the working directory as “host_MDS.png”. See Figure 7.

#### 21. Construct a hierarchical clustering plot to visualize sample groupings

> counts.host.mod <- t(cpm(counts.host))

> dist <- dist(counts.host.mod)

> png(“host_HC.png”)

> plot(hclust(dist), main="Hierarchical Clustering Dendrogram")

> dev.off()

Line 1 converts the counts into Counts Per Million (CPM) and then transposes the resulting matrix. The second command generates the distance matrix between each of the 12 samples, and the third command generates a hierarchical clustering dendrogram from the normalized counts in “counts_host” where the most similar samples occupy closer positions in the tree. The plot is saved to the working directory as “host_HC.png”. See Figure 8.

#### 22. Fit the model

> fit <- lmFit(y, design)

> fit <- contrasts.fit(fit, contrasts)

> fit <- eBayes(fit)

The first two commands estimate expression fold changes and standard errors by fitting a linear model to each gene, using the comparisons defined by the contrast matrix (“contrasts”). The third command applies empirical Bayes smoothing to the standard errors to further weaken any outliers.

#### 23. Print the differentially expressed transcripts for both the 1hpi and 24hpi time-points

> top_1hpi <- topTable(fit, coef = “Host_1hpi”, adjust = “fdr”, number = “Inf”, p.value= 0.05, sort.by = “P”)

> top_24hpi <- topTable(fit, coef = “Host_24hpi”, adjust = “fdr”, number = “Inf”, p.value =0.05, sort.by = “P”)

This command prints all the Differentially Expressed Genes (DEG) with a *p*-value of 0.05 or less after correcting for multiple testing using the False Discovery Rate (Benjamini & Hochberg) method. Additionally, a log fold change (LFC) threshold may be included by adding an “lfc = 2” argument, which would return all DEG with a log fold change in expression greater than two. See Table 1.

#### 24. Annotate the differentially expressed transcript tables with gene symbol,description,and type information

> library(org.Hs.eg.db)

> gene.info <- select(org.Hs.eg.db, key = rownames(top_1hpi), keytype = “ENSEMBL”,

columns = c(“ENSEMBL”, “SYMBOL”, “GENENAME”))

> gene.info <- gene.info[!duplicated(gene.info$ENSEMBL),]

> rownames(gene.info) <- gene.info$ENSEMBL

> identical(rownames(top_1hpi), rownames(gene.info))

> gene.info <- gene.info[, -1]

> host_DEG_table <- cbind(top_1hpi, gene.info)

0ften for downstream applications it is necessary to have the gene name or identifier for each DEG. These commands are derived from the Limma documentation [73] and extract gene annotation information stored in both the org.Hs.eg.db R package and the “host_gene.info” that was previously downloaded to the working directory to annotate the DEG list with gene symbol and gene name. Repeat for the “Host_24hpi” DEG list.

#### 25. Write the annotated differentially expressed transcript table to the local hard drive

> write.table(host_DEG_table, file = “Host_DEG_annotated.csv”, sep = “,”, col.names = NA)

This command writes the DEG list to a comma-separated file. Repeat for the “Ct_24hpi” DEG list.

### Method: Bacteria

#### 26 Build bacteria reference index file and map the unmapped reads (bacterial reads) from HISAT2 to the bacteria reference genome with Bowtie2

$ bowtie2-build -f bacteria_reference.fa bacteria_reference_index

$ bowtie2 -q pair1_unmapped.fastq bacteria_reference_index

The first command indexes the bacteria reference genome “-f bacteria_reference.fa” and generates the “bacteria_reference_index” output file. The second command performs the read mapping using the unmapped reads from the Host mapping step (“pair1_unmapped.fastq”).

#### 27 Repeat steps 6-10 from the host-specific protocol above.

Sort the BAM files by both name and position, convert the “sorted by position” BAM files to BigWig format and visualize with IGV. Create count matrix with HT seq.

#### 28 Repeat steps 13-16 from the host-specific protocol above.

Combine the count files into a DGEList, remove the last five rows from the counts, filter counts to remove low expression genes, and inspect the counts for errors.

#### 29 Apply TMM normalization to counts

> dge.bacteria <- calcNormFactors(dge_bacteria)

> bacteria.cpm <- cpm(dge.bacteria, normalized.lib.sizes = TRUE)

The first command generates TMM normalization factors and the second command converts the raw counts to normalized counts.

#### 30 Calculate gene lengths

> library(GenomicFeatures)

> txdb <- makeTxDbFromGFF(“bacteria_annotation.gtf”, format = “gtf”)

> exons <- exonsBy(txdb, by = “gene”)

> gene.length <- sum(width(reduce(exons)))

> gene.length <- as.data.frame(gene.length)

These commands utilize the the GenomicFeatures R package to extract the gene lengths from the “bacteria_annotation.gtf” file, which are required for the calculation of TPM values (below).

#### 31 Define a function to calculate TPM

> TPM <- function(counts, lengths){

rate<counts/lengths rate/sum(rate)*1e6

}

#### 32 Calculate TPM values

> final.tpm <- apply(bacteria.cpm, 2, function(x) TPM(x, gene.length))

> final.tpm <- as.data.frame(final.tpm)

> colnames(final.tpm) <- colnames(bacteria.cpm)

This command converts the normalized counts in "bacteria.cpm" to TPM.

#### 33 Write TPM values to file

> write.table(final.tpm, file = “Ct_relativeabundance.csv”, sep =“,”, col.names = NA)

## Anticipated Results and Troubleshooting

### RNA quality and quantity

The Bioanalyzer trace will provide an indication of sample quality, where large, well-defined and high molecular weight peaks are expected. Low molecular weight peaks with low definition are usually evidence of RNA degradation and the experiment should not proceed further if this is the case.

### Raw sequence quality checks

Following the analysis of the FASTQ files with FASTQC (step 2), a HTML report is generated which provides a judgement on several sequence quality parameters. 0f particular importance are the following: The per base sequence quality plot should indicate a lower quartile above 10 (corresponding to 90% accuracy), while the per sequence quality score should have a mean base quality of 25 or higher (for RNA-seq reads, it is expected that base quality will decrease in later sequencing cycles), with the histogram curved to the right. The per base sequence content plot should indicate an even proportion of each base, and the per sequence GC content plot should demonstrate a normal distribution of GC content; an abnormal distribution is likely evidence of contamination. In both the per base sequence content plot and per sequence GC content plot, a bias will be observed at the first bases which is specific to Illumina sequencing. The k-mer content plot will indicate the presence of k-mers at the early bases (also Illumina sequencing bias), but should otherwise not be at a higher than expected frequency. The read length distribution plot should indicate that the read length is at least 50 nucleotides, as represented by a single peak.

### Visualize alignments in IGV

IGV allows the visualization of reads mapped to the reference genome and provides an efficient way of identifying problematic data. The alignments should agree with the known gene structures such as intron placements, and the reconstructed transcripts should sufficiently represent the alignments. Verify that genes are roughly evenly covered with reads and if there are known differentially expressed genes, confirm differential coverage between sample groups. Splice junctions can also be visualized for the host reads.

### Feature counting

HTseq will generate a separate tab-delimited file for all samples and replicates, containing the read count for every gene in the annotation file. This file is usually represented with two columns: the Ensembl gene identifier, and the read count: a number indicating how many reads overlap with the feature (gene) listed in the annotation file [5].

### Multi-dimensional scaling plot and hierarchical clustering plot

In the MDS plot, replicate samples from each condition should generally cluster together, although some biological variation is expected. High sample variation may indicate noisy data and can be addressed by removing outlier samples, applying sample-specific quality weights as described in step 20, or applying additional normalization factors using specialized packages such as RUVseq [74]. Samples that cluster according to technical parameters, such as the time and date of sample collection, usually indicates evidence of batch effects which can be corrected by incorporating the effect into the design matrix (step 19). See the Limma user guide for more details on this [73].

### Host differential expression analysis

Following gene annotation (step 25), the table of differentially expressed host genes compiles a number of columns containing significant information comprising the output of the host side of the experiment. The first column lists the Ensembl ID for each gene identified as being differentially expressed, while the remaining columns provide the statistical output for each gene, including (1) the log2 fold-change (log FC) measured between experimental conditions, (2) the average log_2_ expression (AveExp) for each gene, (3) the moderated *t*-statistic, which is the ratio of the log_2_ fold-change to its standard error, (4) the *p*-value, (5) the > 0.05 *p*-value adjusted for multiple testing (via the Benjamini & Hochberg method to control the false discovery rate), (6) the B-statistic representing the log-odds that the gene is differentially expressed, (7) and the gene annotation columns that were incorporated in step 25. This table may be sorted by any column, where the log_2_ fold-change and adjusted *p*-values are usually of most interest.

### Bacterial transcript abundance analysis

The TPM values generated in step 35 are representative of the relative expression levels of the bacteria.

